# Control of Mycobacterium tuberculosis Infection in Lungs is Associated with Recruitment of Antigen-Specific Th1 and Th17 cells Co-expressing CXCR3 and CCR6

**DOI:** 10.1101/2020.03.09.981019

**Authors:** Uma Shanmugasundaram, Allison N Bucsan, Shashank R. Ganatra, Chris Ibegbu, Melanie Quezada, Robert V Blair, Xavier Alvarez, Vijayakumar Velu, Deepak Kaushal, Jyothi Rengarajan

## Abstract

*Mycobacterium tuberculosis* (Mtb)-specific T cell responses associated with immune control during asymptomatic latent tuberculosis infection (LTBI) remain poorly understood. Using a non-human primate (NHP) aerosol model, we studied the kinetics, phenotypes and functions of Mtb antigen-specific T cells in peripheral and lung compartments of Mtb-infected asymptomatic rhesus macaques by longitudinally sampling blood and bronchoalveolar lavage (BAL), for up to 24 weeks post-infection. We found significantly higher frequencies of Mtb-specific effector and memory CD4 and CD8 T cells producing IFN-γ in the airways compared to peripheral blood, which were maintained throughout the study period. Moreover, Mtb-specific IL-17+ and IL-17/IFN-γ double-positive T cells were present in the airways but were largely absent in the periphery, suggesting that balanced mucosal Th_1_/Th_17_ responses are associated with LTBI. The majority of Mtb-specific CD4 T cells that homed to the airways expressed the chemokine receptor CXCR3 and co-expressed CCR6. Notably, CXCR3+CD4+ cells were found in granulomatous and non-granulomatous regions of the lung and inversely correlated with Mtb burden. Our findings provide novel insights into antigen-specific T cell responses associated with asymptomatic Mtb infection that are relevant for developing better strategies to control TB.

## Introduction

After close contact with a person with active tuberculosis (TB), only a minority of individuals develops primary TB disease. The majority of individuals successfully control *Mycobacterium tuberculosis* (Mtb) infection in a clinically asymptomatic state termed latent TB infection (LTBI)(1). Individuals with LTBI are defined as having a positive tuberculin skin test (TST) and/or Interferon-gamma Release Assay (IGRA), a normal chest radiograph and the absence of clinical signs and symptoms of disease(2). Latently-infected individuals are generally thought to contain Mtb within granulomatous lesions in the lung without completely eradicating bacteria, although direct evidence for the persistence of Mtb in human LTBI is lacking. Moreover, it is increasingly recognized that clinically asymptomatic individuals likely reflect a spectrum of infection outcomes, ranging from individuals who may have eliminated infection, to those with low levels of replicating or non-replicating persistent bacteria, to individuals who may harbor actively replicating bacteria without exhibiting overt clinical symptoms(3-5). Identifying immune responses associated with asymptomatic Mtb infection states will provide key insights into mechanisms of immune control that protect against progressing to active TB disease.

Antigen-specific T cell responses are critical for immune control of Mtb infection. In response to Mtb infection, the majority of infected people mount robust CD4 T cell responses involving T helper 1 (Th_1_) cytokines such as IFN-γ and TNF-α, which are important for activating macrophages and curtailing Mtb replication in the lung(6, 7). In addition, IL-17 and Th_17_ responses have emerged as important for protective immunity against TB, but mucosal Mtb-specific Th_17_ responses during immune control of LTBI in humans remain poorly studied. In general, the nature and kinetics of Mtb antigen-specific T cell responses in the blood and lung compartments during human LTBI have not been well characterized. This is in part because small animal models do not reproduce key aspects of human LTBI. Moreover, accurately documenting Mtb exposure, initial infection and early events following infection in humans is almost impossible. Thus, studies of Mtb antigen-specific T cells in humans have been largely confined to cross-sectional characterization of peripheral responses in the blood(8-12). While some studies have examined responses in bronchoalveolar lavage (BAL)(13-15), longitudinal studies in humans comparing Mtb antigen-specific T cell responses in blood and lung compartments have been lacking.

Nonhuman primate (NHP) macaque models of Mtb infection recapitulate multiple features of human Mtb infection, including clinically asymptomatic infection and symptomatic active TB disease (16-18) and are attractive for studying immune parameters associated with control of Mtb infection in peripheral blood and lung compartments. We have previously established a model of LTBI in Indian rhesus macaques, in which low-dose aerosol infection with Mtb CDC1551 leads to the development of asymptomatic Mtb infection. Approximately 80% of infected animals remained disease-free for up to 6 months post-infection (19-24) while only ∼20% progressed to active TB disease. In this study, we characterized the nature, magnitude and kinetics of Mtb antigen-specific CD4 and CD8 T cell responses during asymptomatic LTBI in rhesus macaques over the course of ∼24 weeks post-infection, by serially sampling blood and lung compartments in conjunction with intensive clinical monitoring. We found significantly higher frequencies of Mtb-specific effector and memory CD4 and CD8 T cells producing IFN-γ in the airways compared to peripheral blood; and these were maintained throughout the 24-week study period. Moreover, Mtb-specific IL-17+ and IL-17/IFN-γ double-positive T cells were present in the airways but were largely absent in peripheral blood. The majority of Mtb-specific CD4 T cells that homed to the airways expressed the chemokine receptor CXCR3 and co-expressed CCR6. Notably, CXCR3+CD4+ cells were also found in the lungs of animal with LTBI and active TB and were associated with lower Mtb burdens. Our findings provide new insights into antigen-specific T cell responses associated with the establishment and maintenance of asymptomatic infection.

## Results

### Experimental design and clinical characteristics of rhesus macaques with asymptomatic Mtb infection

Six animals with no clinical signs or symptoms of disease were studied over the course of ∼24 weeks following low-dose aerosol infection (**Figure 1A**). These animals had a median chest radiograph (CXR) score of 0.4, denoting no pulmonary lesions, and maintained normal CRP levels **(Figure 1B)**, body weight **(Supplementary Figure 1A**) and temperature **(Supplementary Figure 1B)**. All animals except one had detectable bacteria upon plating BAL **(Figure 1C)** and three of these animals had detectable, albeit low (< 4-logs), lung bacterial loads at necropsy **(Figure 1D)**. Examination of Hematoxylin-Eosin (H&E)-stained lung tissue at necropsy (at ∼24 weeks post-infection) showed that animals harbored varying degrees of inflammatory lesions, ranging from 0.2% to 20% of the lung being constituted by granulomas **(Figure 1E)**. These results highlight the heterogeneity of clinically asymptomatic animals that control Mtb infection, consistent with the idea that asymptomatic LTBI is represented by a spectrum of Mtb infection states(3, 5).

**Figure 1:**
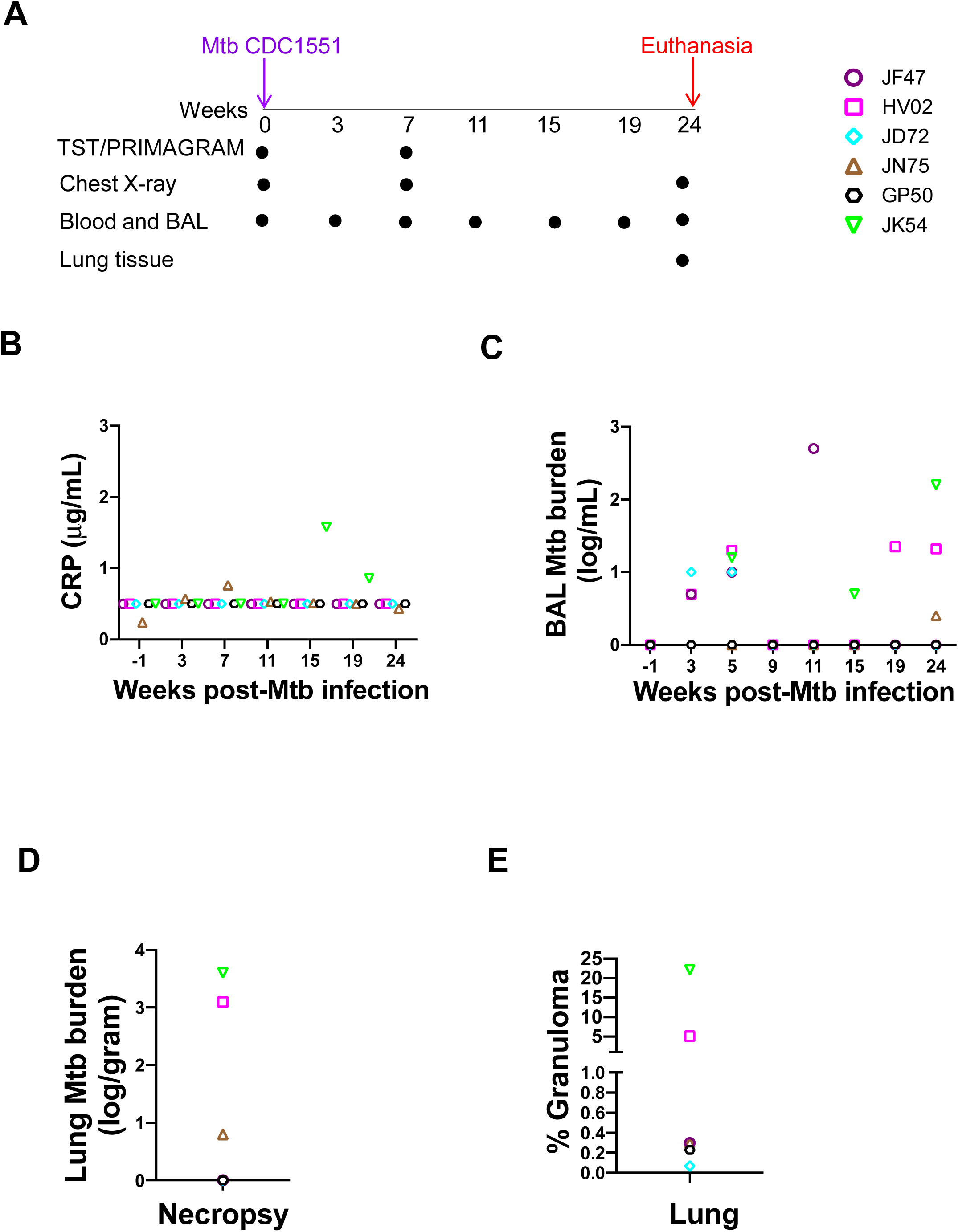
Experimental design and clinical characteristics of asymptomatic rhesus macaques with LTBI. (A) Eight Rhesus macaques were infected by low dose aerosol route with Mtb CDC1551 at week 0 and infection was confirmed at 3 and 7 weeks by TST and IGRA tests. Six macaques were defined as LTBI based absence of clinical signs and symptoms of disease and a negative chest X-ray up to week 15. Six animals remained asymptomatic and were longitudinally followed up until week 24. PBMC and BAL were collected at indicated time points and lung tissues were collected at necropsy. Each colored symbol represents an animal. (B) CRP levels were measured prior to infection (week-1) and at indicated time points post-infection for n=6 animals. (C) Mtb burden (CFU) in BAL measured at indicated weeks (D) Mtb burden (CFU) in lung at necropsy (CFU per gram of tissue plated). (E) Percentage of granuloma in the lung tissue, determined by dividing the granulomatous area (mm^2^) by the area of the annotated regions (mm^2^) classified using an algorithm trained via a deep convolutional network (HALO AI).

### Comparison of total CD4 and CD8 T cell frequencies in PBMC and BAL of asymptomatic rhesus macaques

We assessed the frequencies of total CD4 and CD8 T cells before Mtb infection (week-1) and at 3, 7, 11, 15, 19 and 24 weeks post-infection in the six animals with asymptomatic Mtb infection (**Figure 2**). We observed significantly higher frequencies of CD4 T cells in peripheral blood compared to BAL at all time points studied (*p*=0.01). In contrast, higher frequencies of CD8 T cells were present in BAL compared to peripheral blood at all time points (*p*=0.01). Overall, CD4 and CD8 T cells were each maintained over time in both BAL and PBMCs during the course of the study.

**Figure 2:**
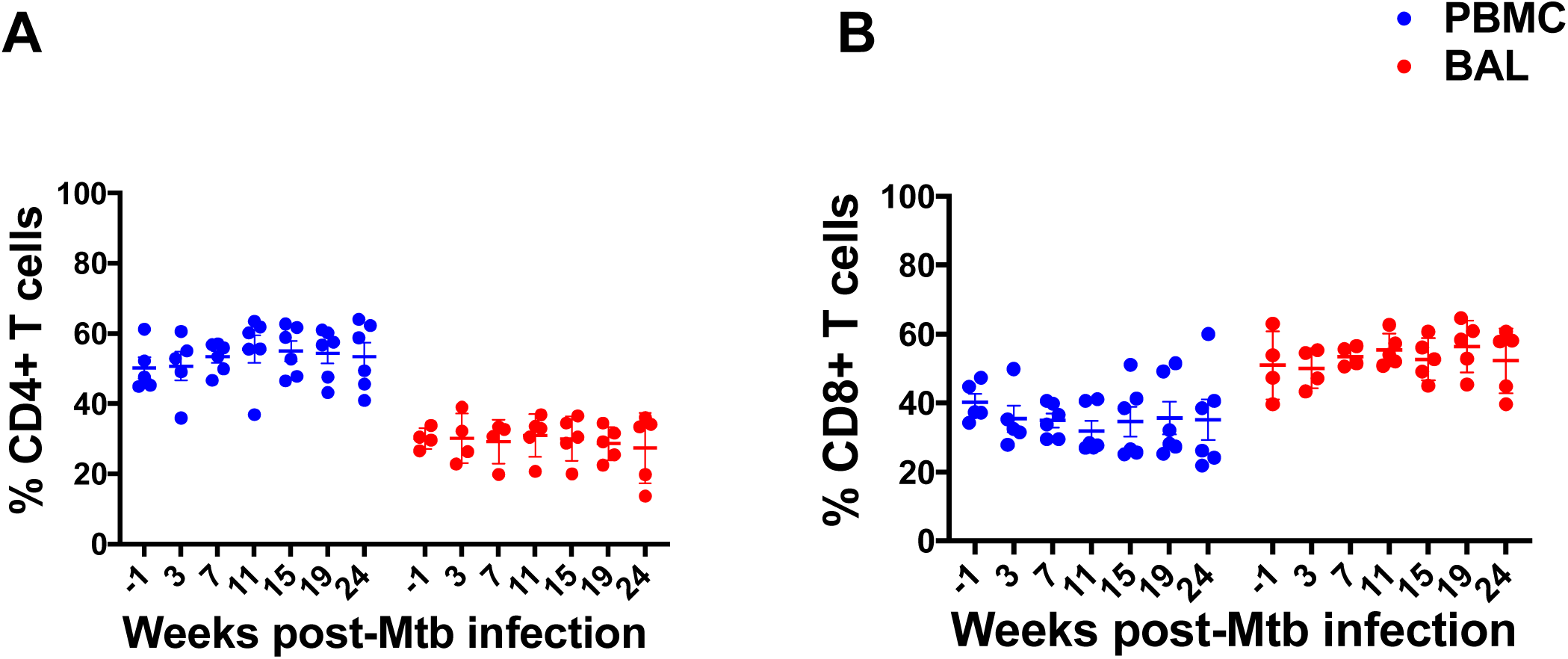
Frequencies of CD4 and CD8 T cells in BAL and PBMC. Red and Blue circles indicate BAL and PBMC respectively. (A) Higher frequencies of CD4 T cells in PBMC compared to BAL at all indicated time points (*p*=0.01). (B) Higher frequencies of CD8 T cells in BAL compared to PBMC at all indicated time points (*p*=0.01). Each data point represents an animal and horizontal lines indicate the mean with SEM. Wilcoxon matched-pairs signed rank test was used to compare the frequencies of CD4 and CD8 T cells between BAL and PBMC.

### High frequencies of Mtb antigen-specific CD4 and CD8 T cells producing IFN-γ are maintained in the BAL of rhesus macaques during asymptomatic Mtb infection

To assess the kinetics of Mtb antigen-specific CD4 and CD8 T cell responses in the peripheral blood and airways of asymptomatic animals that control Mtb infection as LTBI, we stimulated PBMC and BAL samples at each time point with Mtb cell wall antigens (CW) and ESAT-6/CFP-10 peptide pools, followed by intracellular cytokine staining (ICS) and flow cytometry to assess production of IFN-γ. In both PBMC and BAL, CW- and ESAT-6/CFP-10-specific, IFN-γ-producing CD4 **(Figure 3A & B, Supplementary Figure 2A)** and CD8 **(Figure 3C & D, Supplementary Figure 2B)** T cells were detected as early as 3 weeks post-Mtb infection. Mtb-specific CD4 and CD8 T cell frequencies increased at week 7 in both PBMC and BAL and were maintained up to the necropsy endpoint. Overall, while frequencies of Mtb-specific IFN-γ+ CD4 were higher than their CD8 counterparts, both CD4 and CD8 T cell frequencies were significantly (p=0.03) higher in BAL compared to peripheral blood at all time points [mean±SEM, at week 7 post Mtb-infection, 0.3%±0.06 in PBMC and 15%±6.5 in BAL]. Thus, robust Mtb antigen-specific CD4 and CD8 T cell responses are maintained in lung compartments during asymptomatic LTBI.

**Figure 3:**
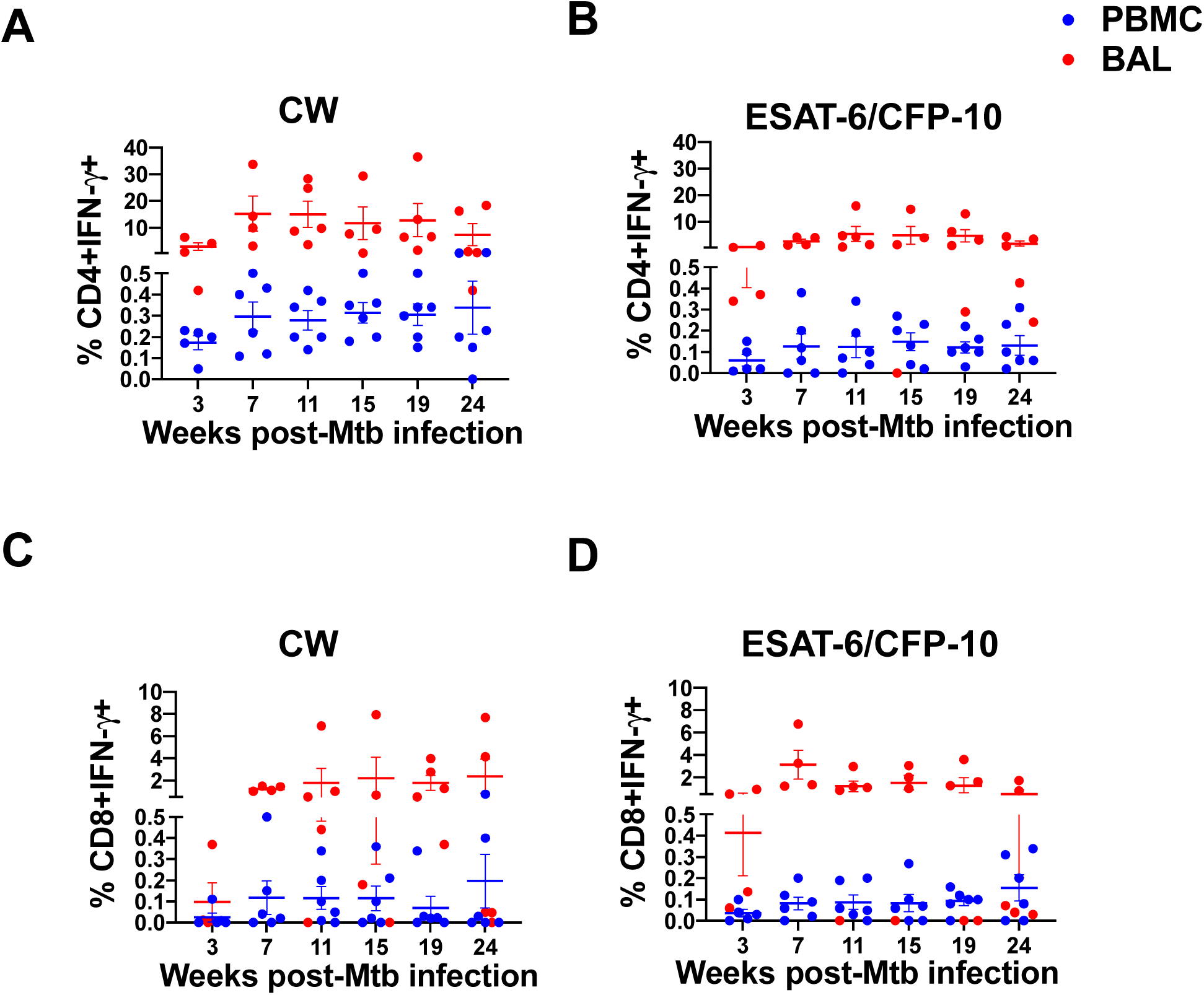
Kinetics of Mtb-specific CD4 and CD8 T cells producing IFN-γ in PBMC and BAL. PBMCs (blue circles) and BAL (red circles) were stimulated with CW and ESAT-6/CFP-10 peptide pools and IFN-γ production by CD4 and CD8 T cells were assessed by ICS and flow cytometry. IFN-γ producing CD4 (A & B) and CD8 (C & D) T cells at indicated time points (weeks) post-Mtb infection in macaques (n=6) with LTBI. CW-specific CD4 (A) and CD8 (C) T cell frequencies and ESAT-6/CFP-10-specific CD4 (B) and CD8 (D) T cell frequencies in PBMC and BAL are shown. Horizontal lines indicate the mean with SEM. Wilcoxon matched-pairs signed rank test was used to compare the frequencies of IFN-γ producing CD4 and CD8 T cells between BAL and PBMC.

### High proportions of CD28+CD95+ Mtb-specific memory T cells in peripheral blood and BAL

The proliferation of antigen-specific T cells in response to infection leads to development of a pool of antigen-experienced memory T cells that are important for mediating effective protection against re-challenge(25). TB vaccine strategies aim to elicit protective antigen-specific memory T cell responses(26, 27). Moreover, memory T cells are thought to play an important role in controlling Mtb infection during LTBI but remain poorly studied in lung compartments. We therefore investigated Mtb-specific effector and memory CD4 and CD8 T cell subsets in asymptomatic rhesus macaques by assessing their differentiation state, based on cell surface expression of CD28 and CD95 on IFN-γ+ CD4 T cells and CD8 T cells **(Figure 4)**. At week 7 post-infection we observed that antigen-specific IFN-γ+ CD4 and CD8 T cells in both PBMC and BAL were predominantly CD28+CD95+ **(Figure 4A-D)**, indicating a central memory-like phenotype. Moreover, these memory T cells were maintained at high levels throughout the time course of the study. While we observed higher proportions of antigen-specific CD28-CD95+ effector CD8 T cells compared to CD4 T cells in the BAL **(Figure 4C, D)**, the relative proportions of CD28+CD95+ and CD28-CD95+ CD4 and CD8 T cell subsets were maintained throughout latent infection in both peripheral blood and airways (**Figure 4E-H**). Our data suggest that antigen-specific memory T cells in the blood and lung compartments are long-lived and are likely to contribute towards maintaining immune control during LTBI.

**Figure 4:**
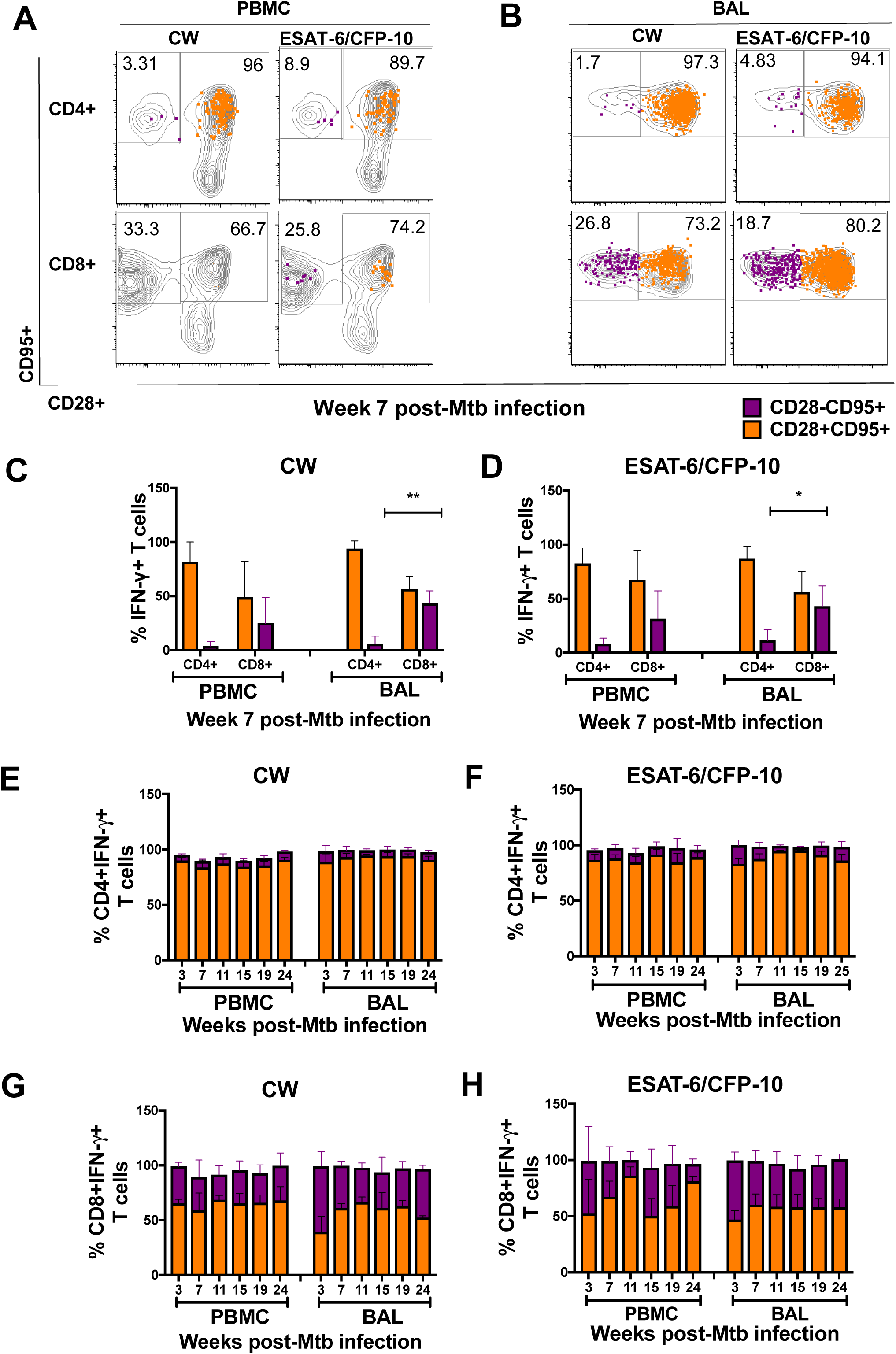
Memory phenotypes of Mtb-specific CD4 and CD8 T cells in PBMC and BAL. (A, B) Representative flow plots of IFN-γ+ CD4 and CD8 T cells expressing CD28 and CD95 in PBMC and in BAL respectively following CW and ESAT-6/CFP-10 stimulation. (C, D) Frequencies of CW-and ESAT-6/CFP-10-specific memory (CD28+CD95+, Purple) and effector (CD28-CD95+, Orange) CD4 and CD8 T cells in PBMC and BAL at week 7 post-Mtb infection. (E-H) Kinetics of CW and ESAT-6/CFP-10 specific memory (Orange bar) and effector (purple bar) CD4 and CD8 T cells in PBMC and BAL. Horizontal lines indicate the mean with SEM. ** *p* <0.01, * *p* <0.05 (paired T-test, 2-tailed).

### IL-17+ and IFN-γ+/IL-17+ T cells are present in the BAL but not in PBMCs from asymptomatic rhesus macaques

In addition to IFN-γ production, IL-17 and Th_17_ responses have emerged as important for protective immunity against TB. To determine whether Mtb-specific IL-17 producing T cells are present during LTBI, we next assessed the kinetics of IL-17+ CD4 and CD8 T cells in both BAL and PBMC **(Figure 5 A-D).** We observed minimal to no Mtb-specific IL-17+ CD4 and CD8 T cell responses in PBMCs throughout the time course of the study **(Figure 5 A-D, Supplementary Figure 2C & D)**. In contrast, CW- and ESAT-6/CFP10-specific IL-17+ CD4 **(Figure 5A & B)** and CD8 cells **(Figure 5C & D)** T cells were clearly detectable in the BAL. Thus, the presence of Mtb-specific IL-17+ Mtb-specific T cells in the airways suggests preferential accumulation of IL-17-producing T cells at lung mucosal sites of Mtb infection and replication.

**Figure 5:**
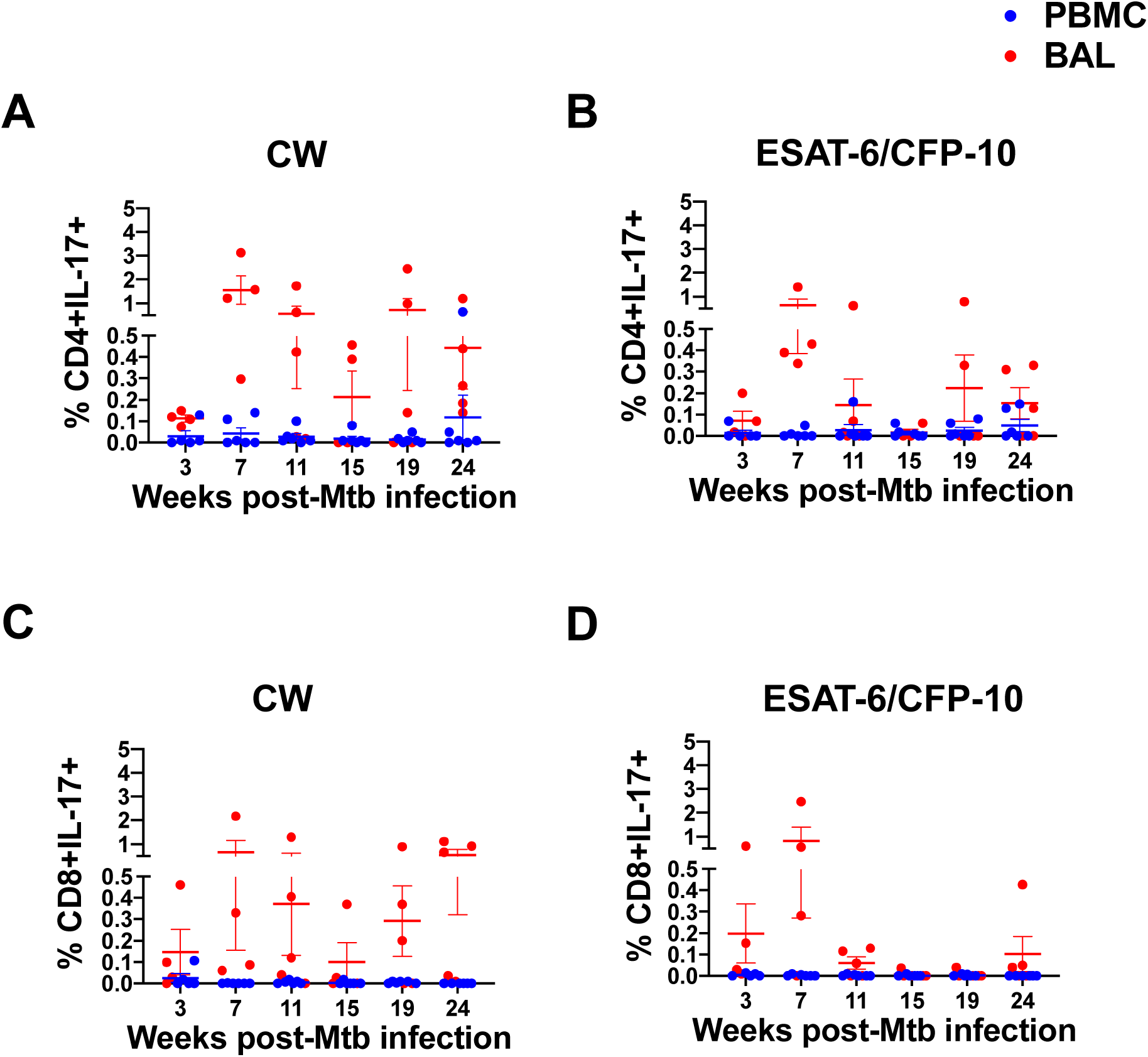
Kinetics of Mtb-specific CD4 and CD8 T cells producing IL-17 in PBMC and BAL. PBMCs (blue circles) and BAL (red circles) samples were stimulated with CW (A & C) and ESAT-6/CFP-10 peptide pools (B & D) and IL-17 production by CD4 (A & B) and CD8 (C & D) T cells assessed by ICS and flow cytometry at the time points indicated (weeks) post-Mtb infection. Horizontal lines indicate the mean with SEM.

Although IL-17 is a hallmark of Th_17_ cells, IL-17/IFN-γ double-positive T cells are also known to be present at mucosal sites of inflammation(28). We sought to investigate whether Mtb-specific CD4 T cells that express both IFN-γ and IL-17 are present in BAL **(Figure 6A).** At week 7 post-Mtb infection, in addition to CW- and ESAT-6/CFP-10-stimulated CD4 T cells that singly-expressed either IFN-γ and IL-17 **(Figure 6A-C)**, we found Mtb-specific T cells that co-expressed IFN-γ and IL-17 **(Figure 6D)**. While these IFN-γ/IL-17 double positive cells were present at relatively low frequencies, our results are consistent with recently published data showing that rhesus macaques who were protected from developing TB following mucosal vaccination with BCG, harbored mycobacteria-specific IFN-γ/IL-17 double-positive CD4 T cells(28). Together, the data represented in **Figures 5 and 6** show that in addition to Mtb antigen-specific IFN-γ+ T cells, Mtb-specific IL-17+ and IFN-γ/IL-17 double-positive T cells are present in the airways of asymptomatic rhesus macaques, suggesting an association between balanced Th_1_/Th_17_ responses at mucosal sites of infection and immune control of Mtb infection.

**Figure 6:**
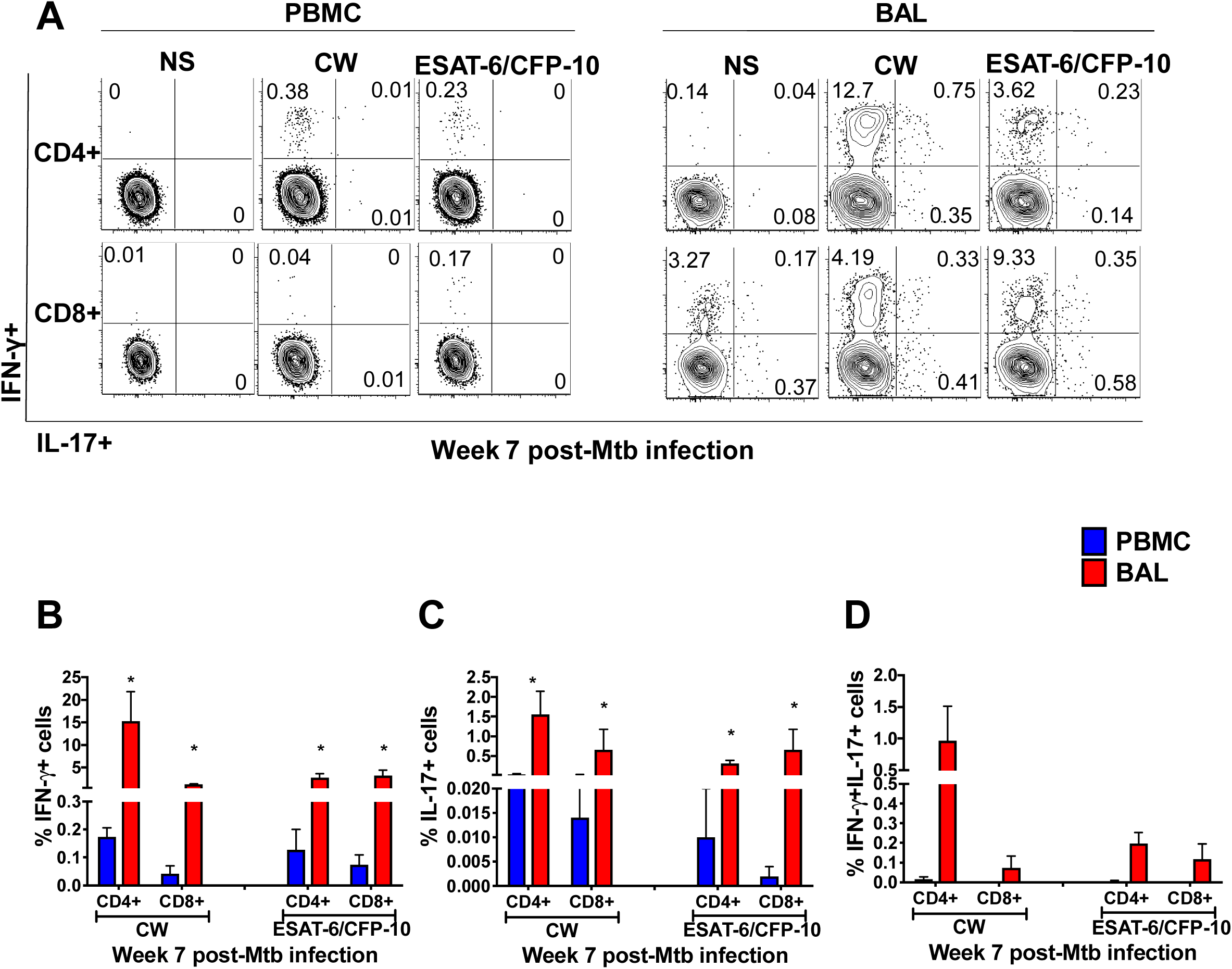
Higher frequencies of IL-17+ and IFN-γ IL-17 double positive Mtb-specific CD4 T cells in the BAL compared to PBMCs. (A) Representative flow plots of week 7 PBMC and BAL that are either non-stimulated (NS) or stimulated with CW- and ESAT-6/CFP-10 peptide pools. CD4 and CD8 T cells expressing IFN-γ and IL-17 were assessed by ICS and flow cytometry. Frequencies of single-positive IFN-γ (B) single-positive IL-17 (C) and IFN-γ/IL-17 double-positive (D) CD4 and CD8 T cells in PBMC (Blue) and BAL (red). \**p* < 0.05 using a nonparametric Mann-Whitney test. Data are represented as mean with SEM.

### Mtb-specific IFN-γ + and IL-17+ CD4 T cells in the airways co-express CXCR3 and CCR6

Chemokine receptors CXCR3 and CCR6 regulate the migration of antigen-specific Th_1_ and Th_17_ cells, respectively, into inflamed mucosal tissues following microbial infection(29-31). We therefore sought to determine whether these chemokine receptors were expressed on the Mtb-specific IFN-γ+ and IL-17+ CD4 T cells present in the airways of macaques that controlled Mtb infection. We assessed the frequencies of CW- and ESAT-6/CFP-10-specific IFN-γ+ and IL-17+ CD4 T cells in the BAL that expressed either CXCR3 or CCR6 **(Figure 7A, B)** and found that the majority of IFN-γ - and IL-17-producing CD4 T cells expressed CXCR3 (mean±SEM at week 7 post-Mtb infection, IFN-γ+ 82.4%±2.9, IL-17+ 61%±7) and CCR6 (mean±SEM at week 7 post Mtb infection, IFN-γ+ 56%±8.6, IL-17+ 67%±11). To assess co-expression of CXCR3 and CCR6, we analyzed samples from the week 7-time point, since the low frequencies of IL-17-producing T cells at later time points precluded reliable analyses. We found that the majority of Mtb-specific IFN-γ + **(Figure 7C, D)** and IL-17+ **(Figure 7E, F)** CD4 T cells in BAL co-expressed CXCR3 and CCR6. Moreover, IFN-γ/IL-17 double-positive cells were also predominantly CXCR3+CCR6+ **(Figure 7G, H).** Overall, CD4 T cells co-expressing CXCR3 and CCR6 were the main IFN-γ - and IL-17-producing subsets present in the airways of macaques with LTBI.

**Figure 7:**
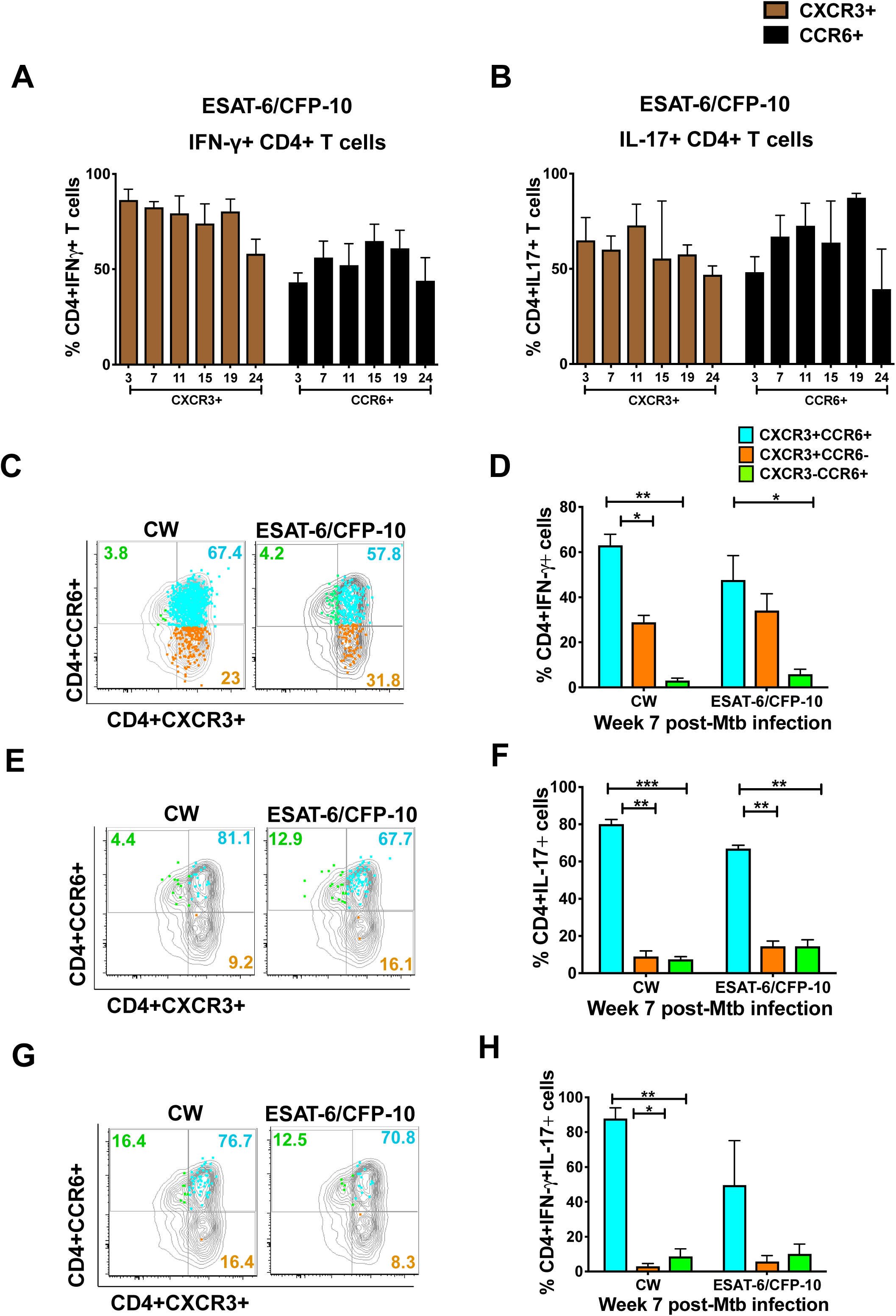
Mtb-specific CD4+ T cells producing IFN-γ and IL-17 co-expressed CXCR3 and CCR6. Frequencies of ESAT-6/CFP-10-specific IFN-γ+ (A) and IL-17+ (B) CD4 T cells in the BAL that expressed either CXCR3 (brown bar) or CCR6 (black bar). Majority of IFN-γ and IL-17 producing CD4 T cells expressed CXCR3 at all time points. Mtb-specific IFN-γ+ (C & D) and IL-17+ (E & F) CD4 T cells in BAL co-expressing CXCR3 and CCR6 (Blue bar) were significantly higher compared to CXCR3+CCR6- (orange bar) and CXCR3-CCR6+ (Green bar) subsets. IFN-γ/IL-17 double-positive cells were also predominantly CXCR3+CCR6+ (G & H). Horizontal lines indicate mean ± SEM. *** *p* <0.001, ** *p* <0.01, * *p* <0.05 (paired T test, 2-tailed).

### CXCR3+CD4 T cells in the lung correlate with lung Mtb burden

Our observation that CXCR3 and CCR6 co-expressing cells were the predominant Mtb antigen-specific CD4+ T cells in the airways of macaques with LTBI prompted us to investigate the localization and frequencies of these cells in the lungs of Mtb-infected NHP by immunostaining of lung tissue sections. While CXCR3+ CD4 T cells were clearly detected in lung tissue, robust immunostaining for CCR6 in paraffin-fixed lung tissue could not be established, despite extensive efforts. Since most of the CCR6+ CD4 T cells in BAL were also positive for CXCR3 **(Figure 7)**, we used CXCR3 as a proxy for both chemokine receptors **(Figure 8).** We also stained lung sections with antibodies to CD163 and CD68 to identify macrophages(32). Immunostaining of lung sections from animals with asymptomatic LTBI showed that CXCR3+ CD4 T cells were present in lung tissue **(Figure 8A)**, with higher densities of CXCR3+ CD4 T cells in granulomatous areas of the lung compared to non-granulomatous areas (**Figure 8B**). Next, to investigate the relationship between lung CD4+CXCR3+ cells and Mtb burden, we stained archived lung tissue sections from rhesus macaques with active TB disease(24). Quantification of the density of CXCR3+CD4+ cells in the lungs of animals with active TB showed that the density of CXCR3+CD4+ cells was significantly (p=0.03) higher in the granulomatous areas compared to non-granulomatous areas of the lung, with overall lower CXCR3+CD4+ densities in the lungs of active TB relative to LTBI **(Figure 8B)**. Interestingly, we observed a negative correlation between lung CXCR3+CD4+ densities and lung CFU **(Figure 8C)**, suggesting that CXCR3+ CD4 T cells are associated with lower Mtb burdens. These results, along with the results reported in **Figures 6 and 7**, suggest that recruitment of CXCR3- and CCR6-expressing Th_1_ and Th_17_ subsets to the airways and lungs of macaques contributes to immune control of Mtb burden.

**Figure 8:**
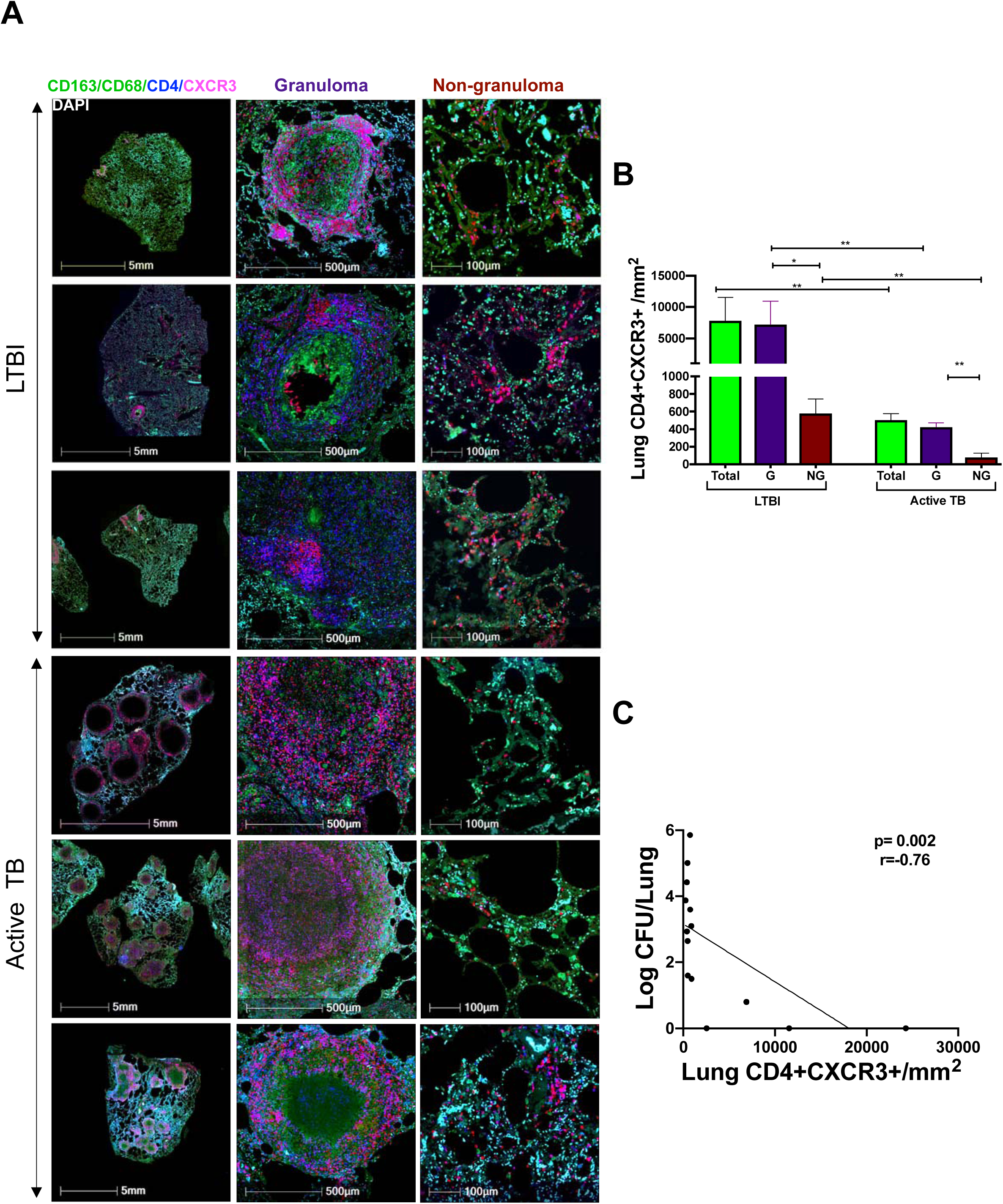
CXCR3+CD4 T cells were predominant in the granulomatous region of the lung. (A) Representative immunohistochemistry staining of lung sections from macaques with LTBI and active TB. Macrophages (green), CD4 and CXCR3 (red) and nuclei (DAPI). Left panel shows low (5mm) magnification images of lung sections from macaques with asymptomatic LTBI and active TB (N=3, each group). Middle and right panels show images of the granuloma (500μm) and non-granuloma (100μm) areas of lung sections, respectively (B) Densities of CD4+CXCR3+ cells in lung tissue of animals with LTBI and active TB in the lung (total) and in granulomatous (G) and non-granulomatous (NG) areas of the lung. Density of CD4+CXCR3+ cells was measured by dividing the number of CD4+CXCR3+ cells quantified by the area in mm^2^ calculated from H&E stained tissue using algorithms trained via a deep convolutional network (HALO, Indica Labs). Comparison within and between groups were performed using nonparametric Wilcoxon matched-pairs signed rank test and Mann-Whitney test respectively. Horizontal lines indicate mean ± SEM. * p<0.05, ** p<0.01. (C) Lung bacterial burden negatively (*p*= 0.002) correlated with the density of lung CXCR3+ CD4 T cells from animals with LTBI and active TB. Correlation was performed using non-parametric Spearman correlation method.

## Discussion

Asymptomatic IGRA+ individuals with LTBI can remain disease-free for decades but current tests cannot determine whether they harbor bacteria or have cleared infection. Thus, defining immune responses associated with establishing and maintaining human LTBI remains challenging. NHPs, e.g., macaques, recapitulate multiple aspects of human Mtb infection and disease progression and are attractive animal models for studying LTBI (16, 33-38). We recently showed that rhesus macaques with asymptomatic Mtb infection harbor viable Mtb bacteria in their lungs for up to nine months(20). Moreover, co-infection with simian immunodeficiency virus (SIV) induced reactivation to TB disease while treatment with isoniazid and rifapentine for 3 months prevented SIV-mediated reactivation to TB. In the current study, we used our established low-dose aerosol infection model of LTBI to study the nature and kinetics of Mtb antigen-specific T cell responses in six rhesus macaques that remained free of clinical signs and symptoms of TB disease for upto 24 weeks post-infection. We observed varying degrees of lung inflammation and granulomas across all the asymptomatic animals, **(Figure 1E and Figure 8)**, consistent with the heterogeneity reported in cynomolgus macaques with LTBI, where both sterile and non-sterile granulomas were present within a single host (39).

Previous studies in NHPs assessed antigen-specific immune responses in BAL at early time points macaques in infected macaques (35, 40-42), or during vaccination (43-45) (28). However, in-depth analysis of the phenotypes, functionality and kinetics of Mtb antigen-specific immune responses in blood and BAL during establishment and maintenance of asymptomatic latent infection have been lacking. To our knowledge, our current study, in which we undertook monthly collection of blood and BAL samples starting at week 3 post-infection and up to 24 weeks, is the first to longitudinally profile Mtb-specific (CW and ESAT-6/CFP-10-specific) CD4 and CD8 T cell responses during LTBI and to directly compare peripheral blood and lung compartments over an extended period of asymptomatic infection. We found that the frequencies of Mtb CW- and ESAT-6/CFP-10-specific IFN-γ-CD4 and CD8 T cell responses were significantly higher in BAL (51-fold for CD4 and 30-fold for CD8) compared to PBMCs, with higher proportions of CD8 T cells present in BAL compared to their CD4 T cell counterparts. The presence of high frequencies of Mtb-specific T cells in the airways as early as 3 weeks after Mtb infection, and their persistence throughout the study until 24 weeks, suggests that Mtb-specific T cells accumulate at mucosal sites of infection, which might contribute to control of latent Mtb infection. Lack of IFN-γ+ CD4 T cells have been shown to result in loss of Mtb bacterial control in mice (7, 46) and in macaques(47, 48).

Interestingly, in addition to antigen-specific IFN-γ+ CD4 T cells, we also detected Mtb-specific IL-17+ CD4 T cells in the BAL by 3 weeks post-Mtb infection. Mtb-specific IL-17+ cells peaked at week 7 and were detected mainly in BAL, being largely absent in blood. The accumulation of IL-17 cytokine-producing CD4 T cells in lung compartments is consistent with the known preferential accumulation of Th_17_ cells at mucosal sites during infection(49). Various studies in mice have also suggested that Th_17_ cells play an important role in protective immunity against TB, both in the context of vaccination (50) as well as in the context of disease progression (51, 52). In addition, studies in macaques with active TB disease have suggested an association between granuloma IL-17+ cells and upregulation of genes related to Th_17_ cells and control of Mtb infection (53, 54). Interestingly, in addition to antigen-specific IL17+ CD4 T cells, we also observed (IFN-γ+IL-17+) double-positive CD4 T cells in the BAL. Our results indicate that both IFN-γ single-positive and IFN-γ+IL-17+ double-positive cells emerge soon after Mtb infection and persist for several weeks, thus likely contributing to controlling Mtb infection during LTBI. These data support and extend previous studies in mice (50, 55, 56) and in macaques (28, 53) suggesting that balanced Th_1_ and Th_17_ responses are associated with enhanced immunity to TB. Future studies that selectively deplete IL-17 or IFN-γ + CD4 T cells in the context of LTBI or vaccination in macaques will provide more direct evidence for the role of these cells in LTBI. In addition, determining the balance between Th_1_/Th_17_ responses in animals that fail to control Mtb infection at early time points post-infection will be of great interest. Overall, our study contributes new insights into the kinetics of Mtb antigen-specific Th_1_ and Th_17_ responses in blood and lung compartments during asymptomatic LTBI in over extended periods of time.

Elicitation and maintenance of memory responses has been associated with protective immunity(26, 27). In the present study, Mtb antigen specific memory CD4 and CD8 T cells, both in PBMC and BAL, were maintained at high levels throughout the time course of the study, suggesting that these cells are likely to be long-lived and involved in maintenance of the asymptomatic LTBI state. Although CD4 T cells were the predominant subset responding to Mtb infection, we also detected antigen-specific IFN-γ+ CD8 T cells, which increased in frequency between 3 and 7 weeks in parallel with antigen-specific CD4 T cells. The CD8 T cell responses that we observed in macaques that control infection differ from previous mouse studies in which lung CD8 T cell responses were considerably delayed compared to CD4 T cell responses (57). Additionally, our study shows that Mtb-specific CD8 T cells producing IL-17 are present, although co-production of IFN-γ and IL-17 was reduced relative to the CD4 subsets. Overall, our findings show that Mtb-specific T cells produce both IFN-γ and IL-17 in the BAL at higher frequencies compared to blood and suggest that these responses may be associated with control of Mtb during asymptomatic Mtb infection.

Th_1_ and Th_17_ cells can be identified by the expression of chemokine receptors, specifically CXCR3 and CCR6 are considered surface marker for Th1 and Th17 respectively(58). Within the CXCR3 and CCR6 axis, another subset of cells that co-express both CXCR3 and CCR6 have been identified (59) and these cells produce both cytokines IFN-γ and IL-17. These IFN-γ/IL-17 double-positive CD4 T cells have been shown to play a pathogenic role during autoimmune diseases(60) but their role in protective immunity to infection remains unclear. With respect to Mtb-infection, a recent study showed preferential expansion of CXCR3+CCR6- and reduction in CXCR3-CCR6+ CD4 subsets in individuals with TB-IRIS(61). In latently infected-individuals, CD4 T cells that mainly co-express CXCR3 and CCR6 that produced IFN-γ were reported to be present in peripheral blood but these cells did not express IL-17 (59, 62, 63). However, the co-expression pattern of CXCR3 and CCR6 on Mtb-specific T cells present in BAL during latent infection has not been studied. In the current study, we found that the majority of antigen specific CD4 T cells co-expressed CXCR3 and CCR6 in BAL **(Figure 7)** and interestingly, unlike in blood where IL-17 producing cells were absent **(Figure 5 and 6)**, CXCR3+CCR6+ subsets in macaque BAL were IL-17+ and IFN-γ+IL17+ co-producing cells.

In Mtb-infected macaques, CXCR3^+^ Th_1_ cells have been shown to be efficient in localizing to lung parenchyma during active TB (40) and in mouse models, these cells have been implicated in containing salmonella within granulomas (64). However, the location of CXCR3+ CD4 cells in the lung and their association with Mtb control in rhesus macaque lungs during LTBI have not been previously described. Using immunofluorescence staining and quantification of CXCR3+CD4+ cells in lung tissue sections from asymptomatic LTBI and Mtb active animals, we found that CXCR3+CD4+ cells were located largely in granulomatous areas. Interestingly, the density of CXCR3+CD4+ cells in the granuloma correlated inversely with bacterial load **(Figure 8)**. Although we were unable to effectively stain for CCR6 in the lung, since majority of CCR6+ cells in the BAL also expressed CXCR3 **(Figure 7)**, we conclude that CXCR3+CCR6+ T cells subsets are likely to be associated with mycobacterial control in the lung.

Our studies clearly show that lung compartments of macaques that control Mtb infection are populated with high levels of Mtb-specific Th_1_ and Th_17_ cells. These cells co-express CXCR3 and CCR6 and their presence correlates with lowered Mtb burdens in the lung. Our study also highlights the unique value of the macaque model of LTBI for studying correlates of protective immunity to TB at mucosal sites of infection. Future studies focusing on the cross-talk between innate immunity and the development and maintenance of Mtb-specific T cell responses in the BAL and lung will further extend our understanding of immune control of Mtb infection and advance the development of better vaccines and immune therapeutics for TB.

## Methods

### Infection of animals with Mtb

The experimental design of these studies is described in **Figure 1**. Eight Indian-origin adult rhesus macaques (*Macaca mulatta*) were exposed to ∼10 CFU of Mtb CDC1551 as described previously (19) resulting in all animals being infected, as assessed by the development of positive TSTs and IGRAs. The animals were obtained from the Tulane National Primate Research Center (TNPRC) breeding colony. Prior to the study, the animals were quarantined for 90 days and tested by both Tuberculin Skin Test (TST) and an NHP-specific IGRA (PRIMAGAM, Prionics) (19) to ensure they were not previously exposed to Mtb infection. A custom head-only dynamic inhalation system housed within a class III biological safety cabinet was used for this purpose (19, 24, 27, 65). All animals tested positive by PRIMAGAM and TST at 3 and 7 weeks post-infection. Six out of eight animals did not exhibit any signs or symptoms of active TB and were considered to have LTBI when they remained asymptomatic for 15 weeks. All six animals remained asymptomatic for the duration of the 24-week study. Two out of eight animals developed progressive primary TB disease and exhibited pyrexia, wasting, high serum CRP, and other clinical signs of TB by seven weeks post-Mtb infection and were included in a different study. All animal procedures were approved by the Institutional Animal Care and Use Committee (IACUC) of Tulane University, New Orleans, LA, and were performed in strict accordance with NIH guidelines. Mtb-infected animals were housed under BSL-3 conditions.

### Clinical procedures and sample collection

Procedures for weekly complete blood counts (CBC), chemistry, chest radiographs (CXR) and BAL at week 3 and every four weeks thereafter have been described previously (19, 24, 32, 66). CXRs were scored by veterinary clinicians in a blinded fashion on a subjective scale of 0–4, with a score of 0 denoting normal lung and a score of 4 denoting severe tuberculous pneumonia, as previously described (19). Measurements of C-Reactive Protein (CRP), body weight and temperature were performed as described earlier (19), at week-1 and at weeks 3, 7, 11, 15, 19 and 24. Peripheral blood and BAL samples were collected at 3, 7, 11, 15, 19 and 24 weeks after Mtb infection. Peripheral blood mononuclear cells (PBMCs) were isolated from blood and cryopreserved for subsequent antigen-specific flow cytometric assays. BAL samples were obtained by bronchoscopy as previously described (19, 67), using two washes of 40 ml sterile saline, and used for antigen-specific assays and to measure CFUs. Bacterial burden associated with Mtb infection was determined in BAL and in lung at necropsy by plating BAL or homogenized tissue sections as previously described (24, 27, 68, 69). Individual lung lobes were cut into 2-mm thick slabs and stereologically selected for analysis which allows for unbiased selection of lung tissue (70). Approximately 50% of the lung tissue was pooled by lung lobe (n=5/animal), homogenized, serially diluted, and plated in quadruplicate for quantification of bacterial load by CFU. Approximately 30% of the lung tissue was fixed for histological analysis and the remaining tissue was fixed for use in immunohistochemistry and immunofluorescence microscopy. The extent of morphometric lung pathology and the involvement of lung in granulomatous lesions was also determined at necropsy. Animals were euthanized at 24 weeks or upon signs of disease. Humane endpoints were predefined in the IACUC protocol and applied as needed to reduce discomfort (27). All animals were routinely cared for according to the guidelines prescribed by the National Institutes of Health Guide to Laboratory Animal Care. All procedures were approved by IACUC, and the Institutional Biosafety Committee (IBC). The TNPRC facilities are accredited by the American Association for Accreditation of Laboratory Animal Care and licensed by the US Department of Agriculture.

### Antigen-specific assays and flow cytometry of PBMC and BAL samples

Cell preparation tubes (CPT, BD Biosciences) were used for PBMC isolation (71) and PBMCs were cryopreserved in 90% fetal FBS (Hyclone, South Logan, UT) and 10% dimethyl sulfoxide (Sigma-Aldrich, St. Louis, MO, USA) until subsequent batch-testing by intracellular cytokine staining (ICS) and flow cytometry. For processing of BAL, mononuclear cells were isolated by passing through a 70-μM nylon cell strainer (Becton Dickinson Discovery Labware, Bedford, MA) followed by washing in complete media (RPMI-1640 containing 10% FBS, 2 mM glutamine, 100 IU/ml penicillin, and 100 µg/ml streptomycin, Lonza, Walkersville, MD, USA). Isolated BAL cells (approx.1-2 x10^6^) were stimulated with CW antigen and ESAT-6/CFP-10 peptide pools for 2 hrs. at 37°C, 5% CO_2,_ Brefaldin-A was added and cells were further incubated for 4 hrs. at 37°C, 5% CO_2_. The stimulated BAL cells were cryopreserved using 90% fetal FBS (Hyclone, South Logan, UT) and 10% dimethyl sulfoxide (Sigma-Aldrich, St. Louis, MO, USA) and stored in liquid nitrogen until subsequent batch processing for ICS staining and flow cytometry.

### Intracellular staining and flow cytometry

Cryopreserved PBMCs were thawed (all time points for each animal were thawed on the same day) and rested overnight at 37°C, 5% CO_2_ in 10% complete RPMI. PBMCs (approx. 1–2 × 10^6^) were stimulated with Mtb CW antigens (10 µg/ml; BEI Resources) or ESAT-6/CFP-10 peptide pools (10 µg/ml, 15-mers with 11 amino-acid overlap, Genemed Synthesis Inc., San Antonio, TX, USA) for 2 hrs followed by the addition of Brefeldin A (10 µg/ml) (BD Biosciences, San Diego, CA, USA) after which the cells were further incubated for 16 hrs. ICS and flow cytometry was performed as described below. Cryopreserved stimulated BAL cells were thawed (all time points for each animal were thawed on the same day) and processed for ICS and flow cytometry. Stimulated PBMCs and BAL cells were stained for dead cells using the LIVE/DEAD Fixable Near-IR Dead Cell Stain (Life Technologies, OR) and then surface-stained with the following antibodies: CD8-V500 (clone SK1), CCR6-BV711(clone 11A9), CD95-PETR (clone DX2) from BD Biosciences, CD38-FITC (clone AT-1, Stemcell Technologies), CD28-PE-Cy7 (clone CD28.2) and CXCR3-BV605 (clone G025H7) from Biolegend. Cells were permeabilized with Cytofix/Cytoperm Kit (BD Biosciences) and stained intracellularly with CD3-PerCP (clone SP34-2), IFN-γ-Alexa Fluor 700 (clone B27) and IL-17-APC (clone MQ1-17H12) from BD Biosciences and CD4 BV650 (clone OKT4, Biolegend). Cell were fixed with 1% paraformaldehyde before acquisition in an LSR-II flow cytometer (BD Biosciences). Flow cytometry data were analyzed using FlowJo software V10 (Tree Star Inc., San Carlos, CA, USA).

### Histology and quantification of granulomatous areas in the lung

Lung tissues were fixed in zinc-buffered formalin, processed routinely, and stained with hematoxylin and eosin (H&E). The stained tissue sections were digitally scanned with a digital slide scanner (Axio Scan.Z1, Carl Zeiss) and analyzed with computer software (Tissue Classifier, HALO, Indica Labs). Annotation regions were drawn around each tissue section on the slide. Annotated regions were than classified using an algorithm trained via a deep convolutional network (HALO AI) to identify granulomas. All analyses were reviewed by a board-certified veterinary pathologist to confirm their accuracy. Percentage of granuloma within tissue sections was determined by dividing the granuloma area (mm^2^) by the area of the annotated regions (mm^2^).

### Immunohistochemistry and confocal microscopy

In addition to tissue sections from the six asymptomatic animals, we also included archived formalin-fixed, paraffin-embedded lung tissues from animals with active TB that were reported in our previous studies(24) and performed immunostaining and confocal microscopy as previously described(27). Briefly, 5µm tissues sections were mounted on Superfrost Plus Microscope slides, baked overnight at 56°C and passed through Xylene, graded ethanol, and double distilled water to remove paraffin and rehydrate tissue sections. A microwave was used for heat induced epitope retrieval (HIER). Slides were boiled for 20 minutes in a Tris based solution, pH 9 (Vector Labs H-3301), containing 0.01% Tween20. Slides were briefly rinsed in hot, distilled water and transferred to a hot citrate based solution, pH 6.0 (Vector Labs H-3300) where they were allowed to cool to room temperature. Once cool, slides were rinsed in tris buffered saline (TBS) and placed in a black, humidifying chamber where they were incubated with Background Punisher (Biocare Medical BP974H) for 10 minutes, washed with TBS containing 0.01% TritonX100 (TBS-TX100) for 5 minutes, followed by a quick rinse in TBS before being returned to the black chamber to be incubated with serum free protein block (Dako X0909) for 20 minutes. The slides were stained with primary antibodies against the following proteins: CD68 [1:20, mouse IgG1 (Dako, Carpinteria, CA)], CD163 [1:20, mouse IgG1 (Leica Biosystems Buffalo Grove, IL)] CXCR3 [1:20 mouse IgG1 (BD Pharmingen, San Jose, CA)], CD4 [1:20, rabbit (Abcam, Cambridge, MA)] and Dapi nuclear stain [1; 20,000 (Invitrogen, Carlsbad, CA)]. The above primary antibodies were detected with the following secondary antibodies from Molecular Probes at a 1:1000 concentration derived from goat: Goat anti-mouse IgG1 488 (green), Goat anti-mouse IgG1 488 (green), Permanent Red, Goat anti-rabbit 647 (far red). Imaging was performed with a Zeiss Axio Slide Scanner (Carl Zeiss, White Plains, NY), and the images were analyzed with computer software (Hiplex fluorescence, HALO, Indica Labs).

### Quantification of CXCR3^+^ cells by immunofluorescence microscopy

Tissue segmentation was first performed using pattern recognition software (Tissue Classifier, HALO, Indica Labs). A random forest classifier was set at a resolution of 7µm/pixel and to detect a minimum object size of 50µm^2^. The classifier was then trained to detect the following tissue classes by providing multiple examples of each tissue class: granuloma and non-granuloma. Annotation regions were drawn around each piece of tissue on a slide and tissue segmentation was performed on the entire piece of tissue **(Supplement Figure 3)**. Computer software (Hiplex fluorescence, HALO, Indica Labs) was used to quantify the following phenotypes: CD4 cells, CD4+CXCR3+ cells, CXCR3+ cells and CD68+/CD163+ macrophages. The software utilized thresholding to both detect cells and set cut off values for expression of each marker/channel. Thresholds were set by real time visual assessment of known positive and negative cells. Analysis and quantification of cellular phenotypes were performed on each tissue segment as defined above.

### Statistics

Statistical analyses were performed using GraphPad Prism (GraphPad Software, La Jolla, CA). The specific tests used are indicated in each of the figure legends.

## Acknowledgements

We thank Toidi Adekambi for help with PBMC processing and optimization of flow cytometry assays and Lakshmi Chennareddi for help with statistical analysis. We also thank present lab members for their helpful suggestions and Jonathan Kevin Sia for comments on the manuscript. We acknowledge the funding support from the following NIH grants: R01AI111943, R01AI123047, R01AI134240, P51OD011133, and P51OD011104.

## Authors contributions

JR, DK and US conceived the studies. US, ANB, SRG, MQ, CI, XA, and RVB performed experiments. US, JR, DK, ANB, CI, XA and VV performed data analyses. US, JR, DK and VV wrote the manuscript.

## Figure Legends

**Supplementary Figure 1:**
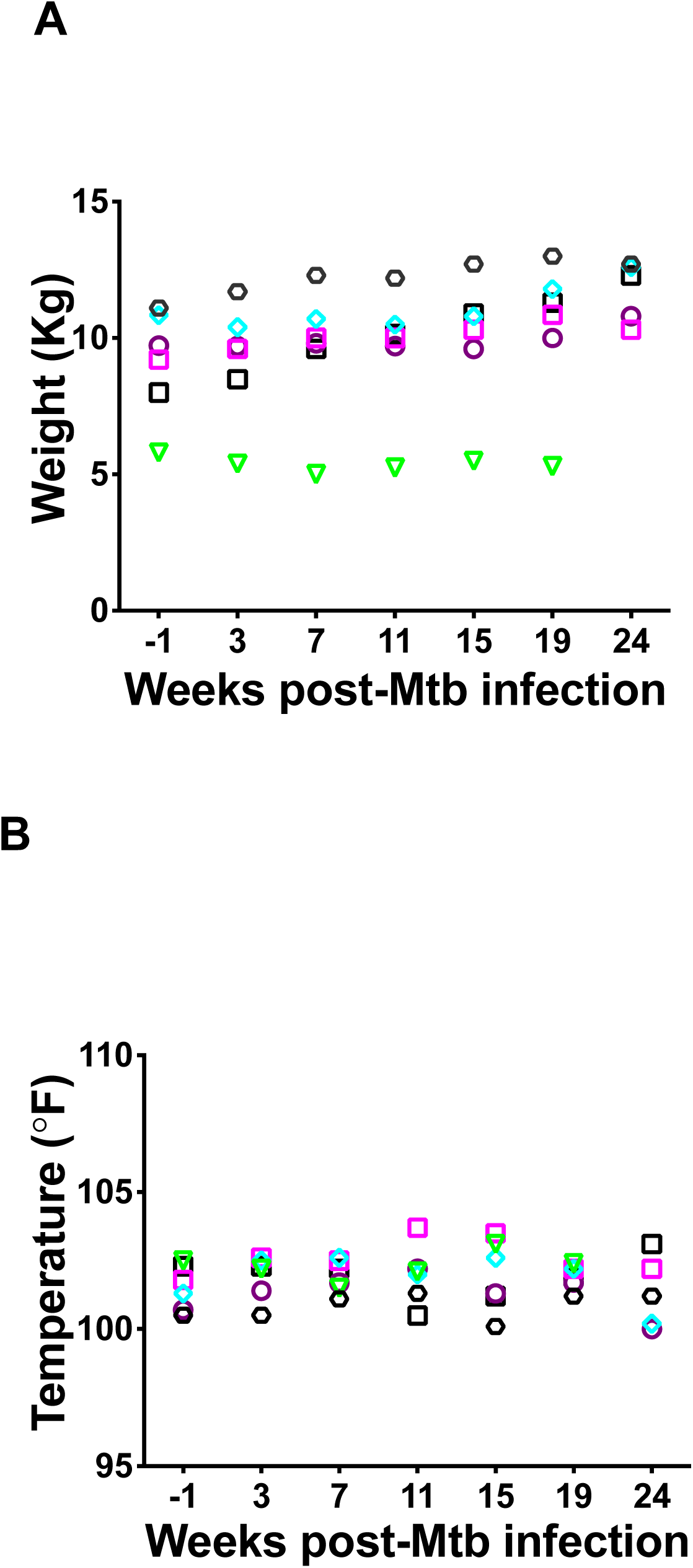
A) Weight and B) Temperature did not change pre- and post-Mtb infection and until week 24 indicating asymptomatic latent infection in rhesus macaques.

**Supplementary Figure 2:**
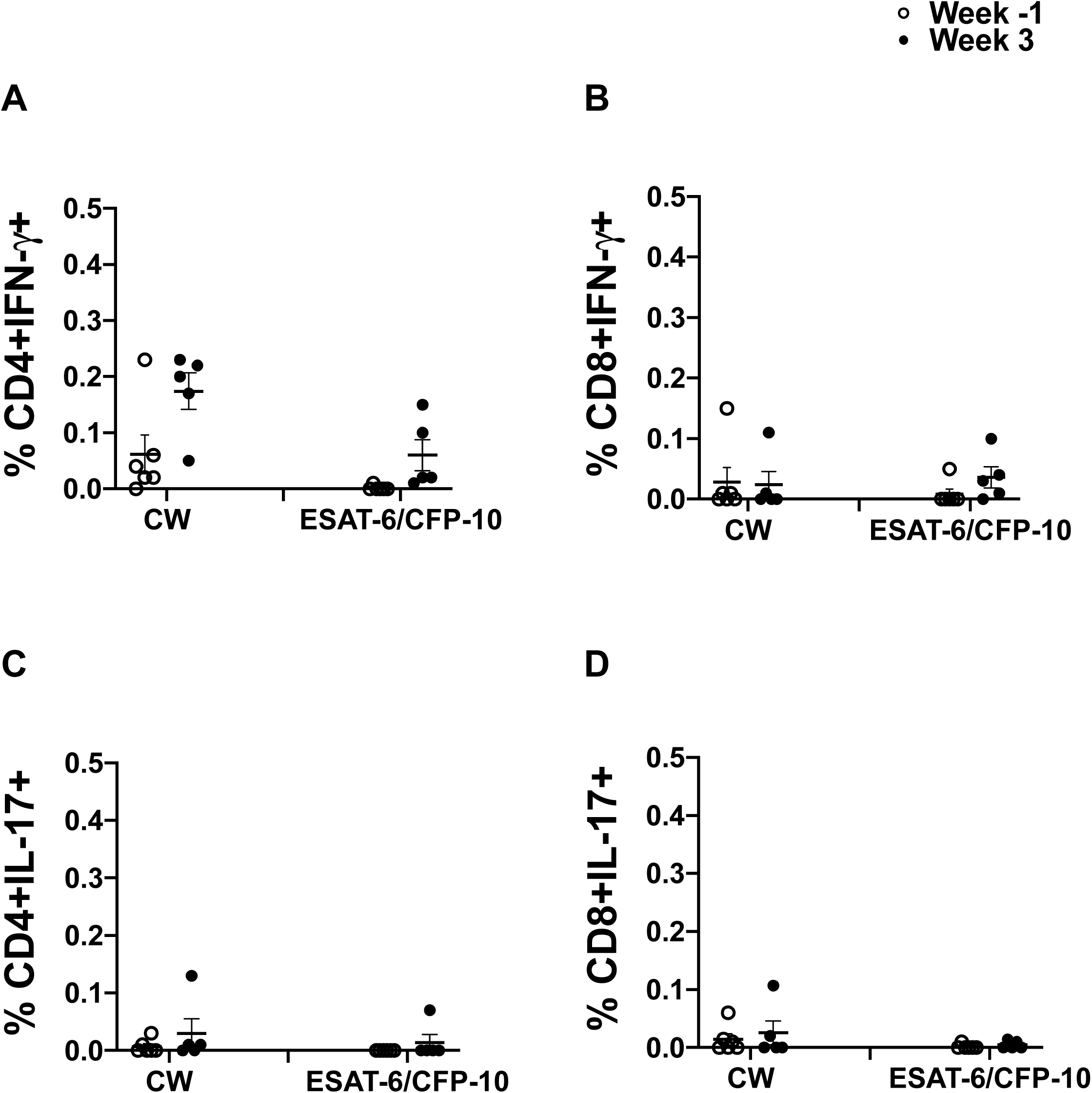
Frequencies of CW- and ESAT-6/CFP-10 specific CD4 and CD8 T cells producing IFN-γ (A & B) and IL-17 (B & D) in PBMC at baseline and week 3.

**Supplementary Figure 3:**
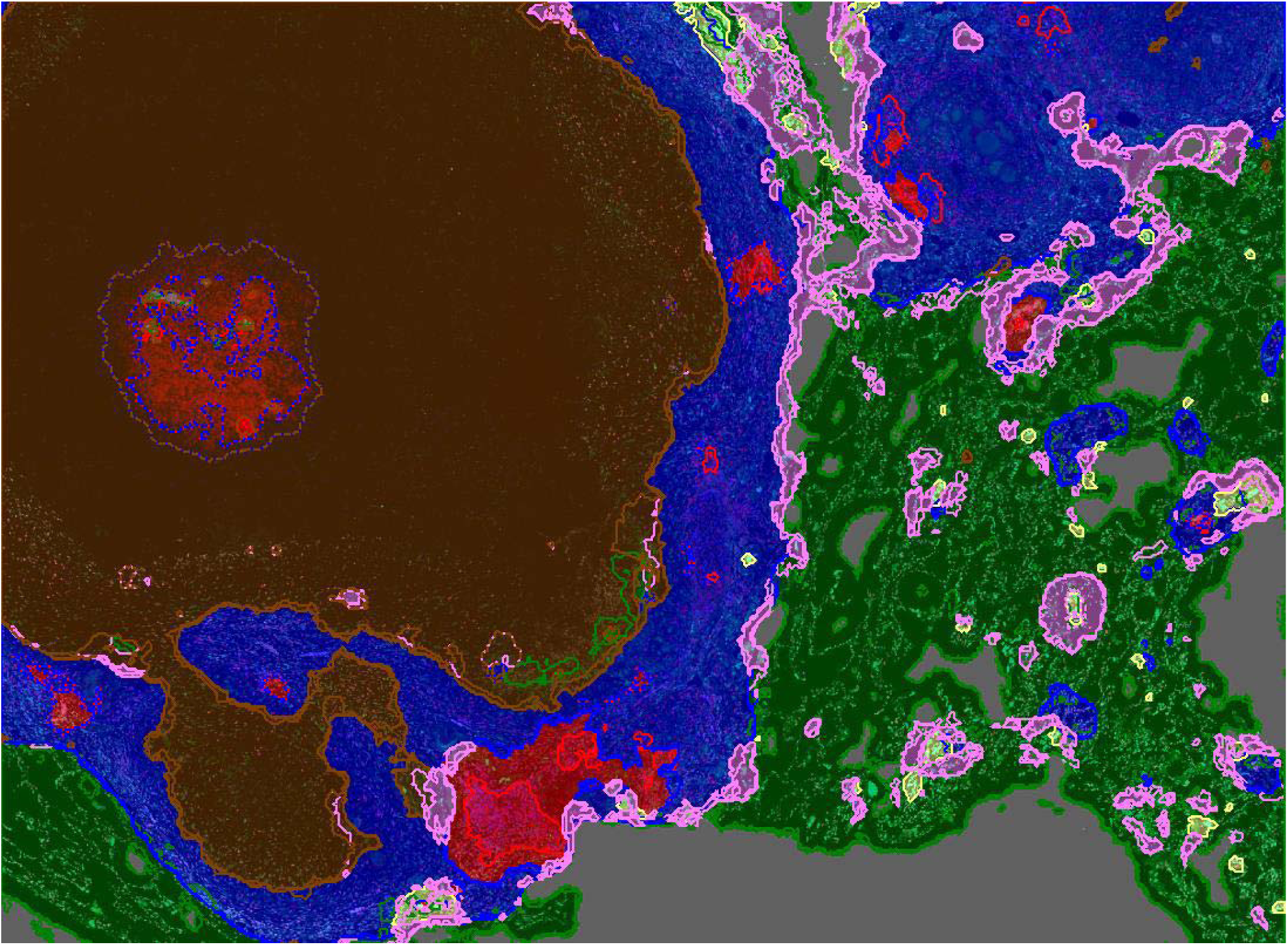
Representative image showing granuloma and non-granuloma region classification: Tissue segmentation performed using pattern recognition software (Tissue Classifier, HALO, Indica Labs). A random forest classifier was set at a resolution of 7um/pixel and to detect a minimum object size of 50μm^2^. The classifier was then trained to detect the following tissue classes by providing multiple examples of each tissue class: granuloma (area in brown and blue) and non-granuloma (area in green). Annotation regions were drawn around each piece of tissue on a slide and tissue segmentation was performed on the entire piece of tissue

